# Varying the position of phospholipid acyl chain unsaturation modulates hopanoid and sterol ordering

**DOI:** 10.1101/2023.09.06.556521

**Authors:** Ha-Ngoc-Anh Nguyen, Liam Sharp, Edward Lyman, James P Saenz

## Abstract

The cell membrane must balance mechanical stability with fluidity to function as both a barrier and an organizational platform. Key to this balance is the thermodynamic ordering of lipids. Most Eukaryotes employ sterols, which are uniquely capable of modulating lipid order to decouple membrane stability from fluidity. Ancient sterol analogues known as hopanoids are found in many bacteria and are proposed as ancestral ordering lipids. The juxtaposition of sterols and hopanoids in extant organisms prompts us to ask why both pathways persist, especially in light of their convergent ability to order lipids. We reveal that both hopanoids and sterols order unsaturated phospholipids differently based on the position of double bonds in the phospholipid’s acyl chain. We find that cholesterol and diplopterol’s methyl group distributions lead to distinct effects on unsaturated lipids. In *Mesoplasma florum*, diplopterol’s constrained ordering capacity reduces membrane resistance to osmotic stress, unlike cholesterol. These findings suggest cholesterol’s broader lipid ordering ability may have facilitated the exploration of a more diverse lipidomic landscape in eukaryotic membranes.

## Introduction

To support life, cell membranes must balance mechanical stability (sufficient to perform robustly as a barrier) against fluidity and deformability (sufficient to support its role as an organizational platform for bioactivity). However, for bilayer forming mixtures of lipids, stability is often gained at the expense of fluidity. Different solutions to this dilemma have evolved on different branches of the tree of life. For example, thermophilic archaea synthesize double headed bolalipids that maintain membrane integrity even at extreme temperatures (1). In many eukaryotes, sterols solve this problem, their twin faces simultaneously ordering hydrocarbon chains while promoting lateral diffusivity (2). By decoupling the local motion of acyl chains and lipid translational freedom of motion, sterols allow cells to build membranes that are mechanically stable enough to withstand environmental perturbations, but fluid enough to support diffusion-dependent biochemistry.

Most non-eukaryotic organisms cannot synthesize sterols and must rely on other mechanisms to modulate their membrane properties. Some bacteria utilize a family of compounds called hopanoids (3). Hopanoids are tri-terpenoids, whose sedimentary record as early as 1.64 billion years ago (4). Since both families are synthesized from squalene, with homologous enzymes (squalene-hopene cyclase and oxidosqualene cyclase) (5), they share certain chemical similarities. Like sterols, hopanoids also reside within the membrane and modulate membrane robustness, fluidity, and resistance against abiotic stresses (3, 6). For these reasons, hopanoids are considered both bacterial and ancient sterol analogs. But why does life need two analogous classes of ordering lipids?

Despite their similarities, hopanoids and sterols possess distinct properties. Both diplopterol (Dpop), a common hopanoid in bacteria, and Chol (Chol), a mammalian sterol, interact favorably with and condense saturated lipids, a diagnostic feature of lipid ordering (7). However, the condensing effect of Dpop is impaired when an acyl chain unsaturation is introduced, and in a manner that depends on the location of the double bond. In this brief report, we examined how double bond position influences sterol and hopanoid ordering. We find that methyl group distribution on hopanoid backbones restricts its ordering ability, resulting in pronounced effects on cellular robustness in the model organism *Mesoplasma florum*. Conversely, Chol ordering is comparatively insensitive to double bond position, allowing eukaryotic membranes more degrees of freedom in regulating their lipidomes.

## Results

Since unsaturation changes the interaction between Chol/Dpop and phospholipids, we hypothesize that moving the position of double bonds along the phospholipid acyl chain can help us probe Chol/Dpop ordering, thus revealing key structural features needed for ordering. We also aim to draw a comparison between Chol and Dpop’s ordering to showcase a constraint in the evolution of sterol- and hopanoid-containing lipidomes.

We first determined the ordering of phospholipid chains by measuring monolayer surface pressure vs area isotherms. Similar isotherms were obtained for PC phospholipids with a double bond at one of three positions: Δ6, Δ9, or Δ11 (chemical structures are shown in Fig. 1A), suggesting similar packing in the pure membranes, regardless of the isomer (Fig 1B). The isotherms differ, however, for binary mixtures of either Chol or Dpop, in a manner dependent on both terpenoid choice and double bond position. To better quantify this difference, we calculated the condensing effect of the terpenoid on each lipid, as well as the free energy of mixing (ΔG_mix_) the phospholipid and terpenoid (see Methods for definitions and details). A positive condensing effect is obtained when the terpenoid orders lipid chains and increases lipid packing, which Chol consistently exhibited regardless of the double bond position. In contrast, Dpop only condenses the Δ11 isomer, does not noticeably affect the Δ6 isomer, and has a negative condensing effect on Δ9 isomer. These observations are mirrored by ΔG_mix_, which reflects the thermodynamic balance between lipid-lipid interactions and mixing entropy. Chol has favorable interaction with all 3 lipid isomers (ΔG_mix_ < 0), which progressively increases as the double bond moves away from the headgroup. On the other hand, Dpop interaction with PC varies based on double bond positions. While Dpop was ideally mixed with Δ6-PC (ΔG_mix_ = 0), the mixing of Dpop and Δ9-PC was unfavorable (ΔG_mix_ >0), and only with Δ11-PC was the ΔG_mix_ <0. The distinction between Dpop interaction with Δ9-PC and Δ11-PC is intriguing, as the double bonds are just 2 carbons apart.

**Fig 1.**
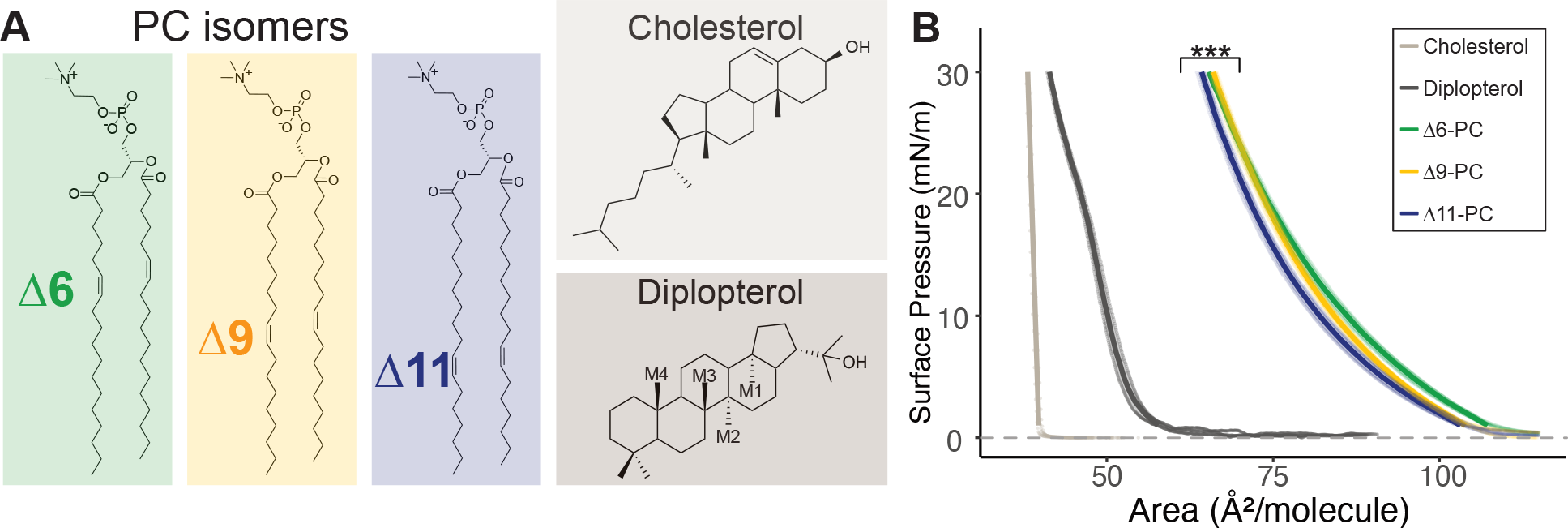
(A) Chemical structure of di-unsaturated PC isomers, Chol and Dpop and (B) Isotherms of the corresponding lipids at 20 °C, *** (F=12.4, p < 0.0005).

**Fig 2.**
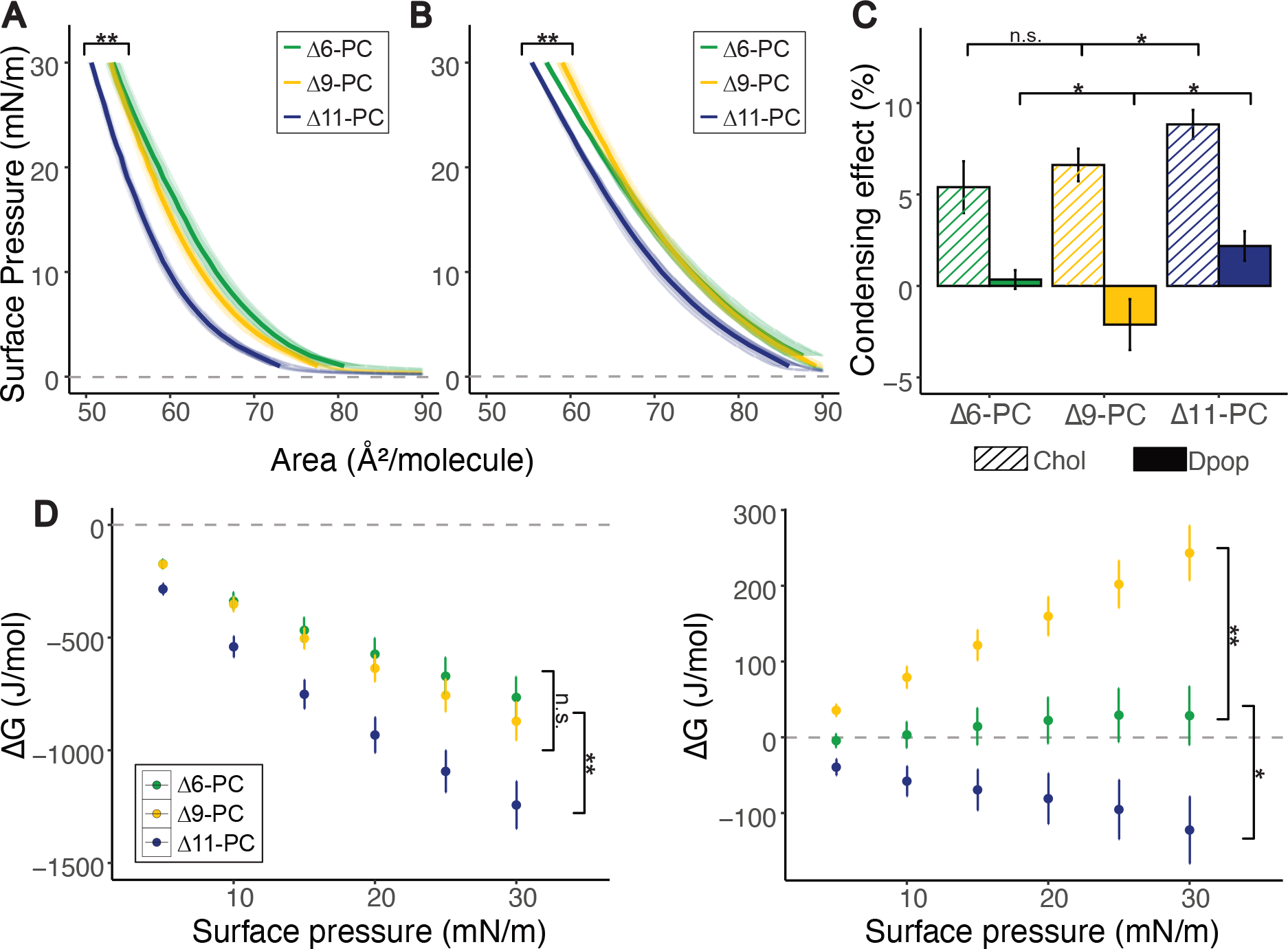
PC isomers interact differently with Chol and Dpop. While Chol ordering of PC increases as double bond position is shifted further from the headgroup, Dpop only exhibits an ordering effect with Δ11-PC. (A) Isotherms of PC isomers mixed with Chol (2:1) at 20°C. ** (F=7.75, p < 0.005) manova (B) Isotherms of PC mixture with Dpop (2:1) at 20°C. ** (F=8.18, p < 0.005) manova (C) Condensing effect of Chol and Dpop on PC isomers calculated at 30 mN/m. Error bar represents standard deviation. n.s. (p > 0.5) * (p < 0.05) unpaired t-test (D) Energy of interaction (ΔG_mix_) of lipid pairs during compression. Error bar represents standard deviation. n.s. (p > 0.5) * (p < 0.05) ** (p < 0.005) unpaired t-test.

To gain molecular insight into differential ordering of Δ9 and Δ11-PC by Dpop we performed molecular dynamics simulations of bilayers of these binary mixtures. (Details in Methods) Figure 3 reports the 2H NMR chain order parameters (S_CD_) for several different binary mixtures of Dpop with phospholipids. S_CD_ is defined in Eq. 2 in methods; larger values indicate more ordered chains. A model for Dpop compatible with the CHARMM36 family of force fields was built as described in Methods, and first tested by simulating binary mixtures with DPPC (Fig. 3A), obtaining an ordering effect similar to that observed in experiments on giant unilamellar vesicles (7).

**Fig 3.**
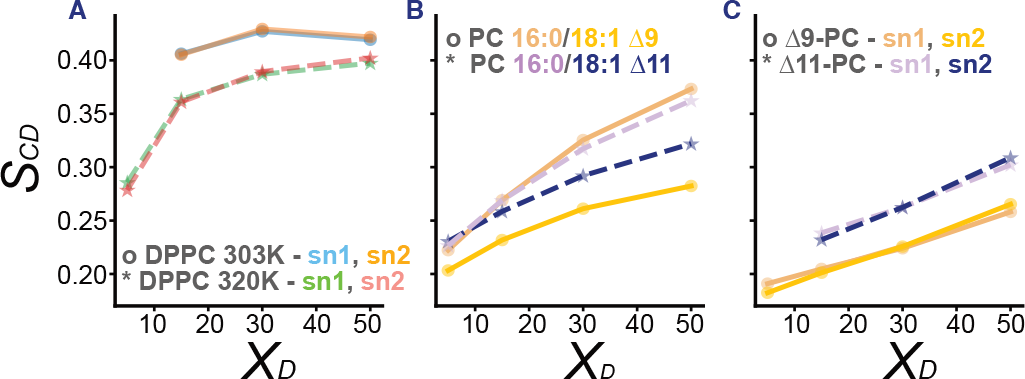
Molecular dynamic simulations show Dpop orders 18:1 Δ11 more efficiently than 18:1 Δ9. (A) Dpop ordering effect with saturated DPPC were confirmed by S_CD_. (B) Dpop ordered saturated chains most efficiently, then 18:1 Δ11 and the least for 18:1 Δ9. (C) Dpop’s ordering effect was similar regardless of sn chain position, but depended on double bond position, with Δ11 a greater ordering effect than Δ9.

We next considered binary mixtures with lipids with saturated chains at the sn-1 position and monounsaturated chains (Δ19 or Δ11 isomers) at the sn-2 position (Fig 3B). As expected, the saturated chains were more ordered by Dpop than the unsaturated chains, and were ordered similarly regardless of which isomer was present in the other chain. Comparing the unsaturated chains, the Δ11 isomer was significantly more ordered than the Δ9 isomer. This trend was reproduced in PC with unsaturation in both acyl chains (Fig. 3C), where we again observed that the Δ11 isomer was significantly more ordered. In summary, the position of the double bond was critical in determining the acyl chain’s order when interacting with Dpop, mirroring the observations in the monolayer experiments and supporting a model in which unsaturation position has a significant effect on condensation in fluid lipid bilayers.

In prior work, Martinez-Seara et al. found that the positioning of the double bond relative to the methyls protruding from cholesterol controls its condensing effect (8). We therefore performed a similar analysis, comparing the location of the PC lipid’s double bond to the positions of methyl groups extending from the Dpop ring structure (annotated in Fig 1A) in our simulations. Figure 4 reports the distribution of double bonds of the hydrocarbon chains and Dpop methyl groups along the membrane normal, with z = 0 being the center of the bilayer. The distribution of Δ9-PC’s double bond overlaps almost exactly with that of methyl group M2, while the Δ11-PC double bond falls in between the M3 and M4 methyl groups. The alignment between the π bond and the methyl group might explain the reduced condensing effect observed for Δ9-PC, while Dpop and Δ11-PC interacted more favorably (8). This interplay between lipid structure and membrane biophysics provides insights that could be extended to other sterols and hopanoids, possibly laying a path for predicting how hopanoid/sterol and phospholipid structure collectively influence membrane properties.

**Fig 4.**
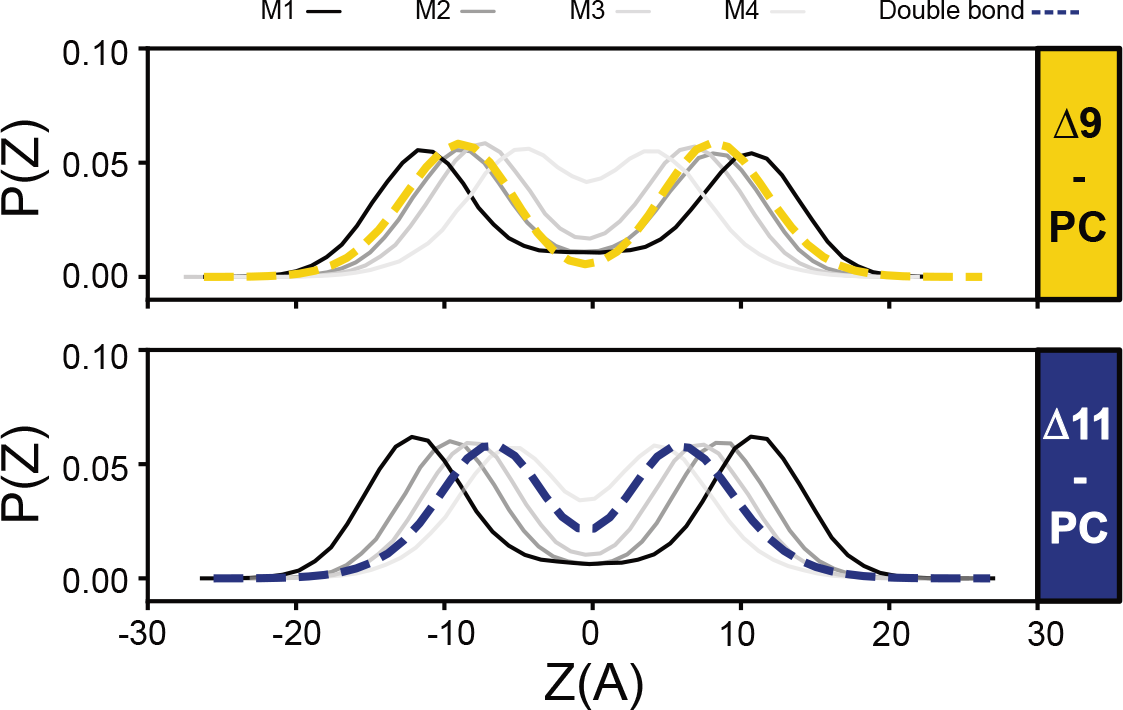
Overlapping distribution of Dpop’s methyl groups and PC’s double bond correspond to a reduced ordering effect. (A) The double bond in Δ9-PC overlaps with Dpop M2, preventing efficient lipid packing. (B) No Dpop methyl group overlaps with the double bond of Δ11-PC.

To investigate how this variation in lipid–lipid interaction might influence biomembrane function, we employed *Mesoplasma florum* as a living model system. *M. florum* is a Mollicute with no cell wall and a minimal genome (9). With limited machinery, Mesoplasma cannot synthesize its own lipids and relies on supplemented lipids from the media, offering a straightforward way to manipulate its lipidomes. By introducing either Δ9- or Δ11-PC to its lipid diet, we can create two identical biological membrane systems differing only in their unsaturation site. We then investigated this system to explore the effect of lipid–lipid interactions on a cellular scale.

Traditionally cultured with an undefined lipid diet in serums, we first test Mesoplasma’s ability to grow in a defined lipid diet. Figures 5A and 5B report growth rate and cellular lipid content, respectively, suggesting Mesoplasma can both grow on and incorporate defined lipids into their membrane. Since sterol ordering has previously been associated with membrane robustness (2), we tested Mesoplasma membrane robustness with hypoosmotic shock. Live cells are subjected to hypoosmotic conditions, forcing cells to rapidly expand. As the membrane is stressed and ruptures, exposed cellular DNA is stained by propidium iodide. By quantifying the fluorescence intensity, we estimated the fraction of cells lysed due to osmotic shock, inferring membrane robustness. Figure 5C shows the cell’s susceptibility to osmotic shock when supplied with different lipid diets. When cells were supplied with Chol, the addition of Δ9- or Δ11-PC did not produce significant changes in cellular robustness. However, cells fed with Δ9-PC and Dpop exhibited higher susceptibility to lysis than cells fed with Δ11-PC and Dpop. This data suggested that Dpop and Δ9-PC unfavorable interaction counteracts Dpop’s ability to bolster membrane robustness.

**Fig 5.**
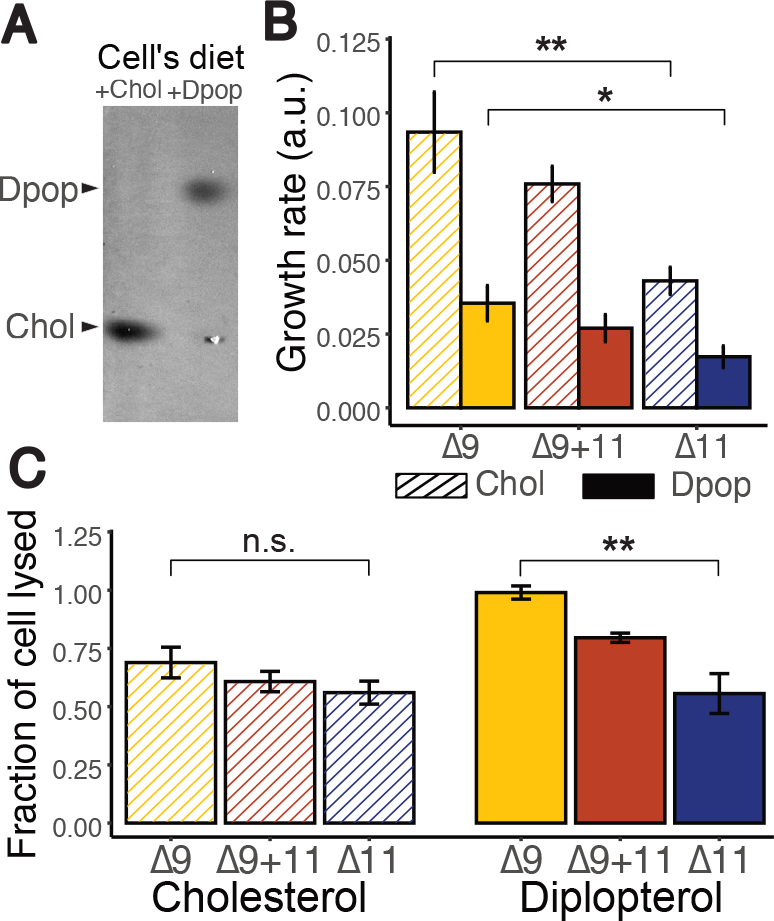
Dpop and Δ11-PC enhance robustness of *Mesoplasma florum* to hypoosmotic shock compared with Dpop and Δ9-PC. (A) Chol and Dpop were incorporated into Mesoplasma membranes according to their respective diets (B) Growth rate of *M. florum* on different diets. ** (F=24.4, p < 0.005) * (F=10.6, p < 0.05) Analyses were performed using one-sided anova with Tukey post hoc test (C) Membrane robustness reflected by the fraction of cell lysed when subjected to hypoosmotic shock. n.s. F=1.47, p<0.5) ** (F=16.6, p < 0.005) Analyses were performed using one-sided anova with Tukey post hoc test.

## Discussion

Chol and Dpop are both prevalent lipids, accounting for approximately 40% of their respective membrane lipidomes (10, 11). We show that both of these tri-terpenoids interact with unsaturated lipids in a double bond position-dependent manner. While Chol condenses all unsaturated isomers comparably, Dpop shows the strongest condensing effect on Δ11-PC, albeit less potently than Chol. Simulations indicate that Dpop’s interaction with unsaturated lipids is significantly hindered by having multiple methyl groups extending from both sides of the cyclic ring structure, similar to the biosynthetic precursor to Chol, lanosterol (12–14). Notably, when the double bond resides at the Δ9 position, and overlaps with Dpop’s methyl group M2, it prevents effective Dpop-induced lipid packing. In combination with earlier simulation results for cholesterol from Martinez-Seara et al. (8), our observations highlight the power of using simulations to explore lipid-lipid interactions, especially in the case of less commercially available lipids like hopanoids. With careful consideration in model development, one can explore the chemical landscape of lipids and the consequences of their collective interactions.

From these observations, we hypothesized that the differentiation between Δ9 and Δ11 should be the most significant in hopanoid-bearing membranes. In *Mesoplasma florum*, the favorable interaction of Δ11-PC and Dpop enhanced membrane resilience to osmotic shock compared to Δ9-PC. This remarkable result highlights how subtle changes to lipid structure can have striking consequences for cellular robustness, and suggests a potential mechanism for osmoadaptation, for which hopanoids have been shown to play a critical role in soil and plant associated bacteria (15, 16). Indeed, multiple hopanoid-bearing bacteria have Δ11 as the monounsaturation site (17), instead of Δ9 in eukaryotes. The hopanoid producing yeast, *Schizosaccharomyces japonicus*, also possess a Δ12-desaturase (18, 19). Interestingly, in 2020, Chwastek et al. investigated the lipidome of a hopanoid-bearing organism *Methylobacterium extorquens*, and found that the main unsaturation position was Δ11 instead of Δ9, with an additional Δ5 unsaturation upregulated in cold-adapted lipidome (11). Therefore, double bond position could represent a modifiable lipidomic feature that cells can employ to homeostatically fine-tune the ordering effects of hopanoids.

For Chol-bearing organisms, double bond position does not have such a pronounced effect on lipid ordering. This indifference might have alleviated evolutionary selection against Δ9 unsaturation in early sterol-bearing organisms, providing more flexibility to produce lipids with double bond positions optimized for orthogonal lipid-lipid or lipid-protein interactions. Our results, therefore, suggest that a transition from hopanoid to sterol-containing lipidomes could have widened the chemical landscape available for cells to explore for tuning membrane properties.

Acknowledgments

The authors thank Tomasz Czerniak, Lisa Junghans, Nataliya Safronova, Isaac Justice and Fabian Rost for helpful discussions and feedback. This work was supported by the B CUBE of the Technische Universität Dresden, a Maria Reich Fellowship (to H.N.A.N), a German Federal Ministry of Education and Research (Bundesministerium für Bildung und Forschung - BMBF) grant (to J.P.S., Project No. 03Z22EN12), and a Volkswagen Foundation ‘‘Life?’’ grant (to J.P.S., Project No. 93090). E.L. and L.S. were supported by NIH award RO1GM120351. This work used Expanse2 at the San Diego Supercomputing Center through allocation BIO230093 from the Advanced Cyberinfrastructure Coordination Ecosystem: Services & Support (ACCESS) program, which is supported by National Science Foundation grants #2138259, #2138286, #2138307, #2137603, and #2138296.

## Materials

Δ6-, Δ9-, Δ11-PC and egg sphingomyelin were purchased from Avanti Polar Lipids. Chol and palmitic acid were purchased from Sigma, and Dpop from Chiron. Stock concentrations of lipids were measured by phosphate assay. Chol and Dpop were weighed out on a precision scale and solubilized in a known volume of chloroform.

## Methods

### Monolayer

Chloroform solutions of pure lipids and mixtures were prepared at 0.2 mg/mL lipid concentrations. Monolayers were prepared by injecting 15-30 μL of lipid solution onto an aqueous subphase maintained at 20°C by a built-in temperature-controlled circulating water bath. The subphase was comprised of 10 mM HEPES, 150 mM NaCl, pH 7. Isotherms were recorded using a 70 cm^2^ teflon Langmuir trough fitted with a motorized compression barrier equipped with pressure sensor (Kibron DeltaPi).

The mean molecular area (MMAs) for each mixture were estimated from the averages of isotherms from three monolayers that were prepared independently. Data were rounded down to the nearest neighbor for condensation effect and free energy calculation. All isotherms were fitted to a regression, and statistical significance was tested using manova with the 2 coefficients. The condensation effect was calculated as follows:

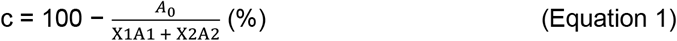

Where c = % condensation, A_o_ = the MMA of the lipid mixture, X_1_, X_2_ = the mole fraction of lipid 1 and 2 in the mix, and A_1_, A_2_ = the MMAs of lipid 1 and 2 at surface pressures 30 mN/m. Error bars were produced based on error propangation.

The ΔG was calculated by integrating the areas of lipid mixtures over pressures Π = 5, 10, 15, 20, and 25 mN/m according to Grzybek et al (20). Error bars were produced based on error propagation.

#### Diplopterol model development

A CHARMM compatible model for diplopterol (Dpop) was developed using the automated atom typing and parameter assignment pipeline CGenFF.30 Charmm topology and parameter files are provided as Supplemental Material.

#### Simulation composition and construction

Simulation systems contained either DOPC or POPC (unsaturation at either the Δ9 or Δ11 position) and one of either Dpop or Chol. All initial configurations were built using the CHARMM-GUI webserver (21–23). Systems containing atypical unsaturated chains (ie, Δ11) were generated by first building a binary mixture of the corresponding Δ9 lipid (DOPC or POPC) with either Dpop or Chol, then “mutating” the unsaturated chain(s) to move the double bond to the appropriate position, using a Charmm script provided by the Klauda Lab.

All simulations contained approximately 550 lipids per leaflet and at least 50 TIP3P (24) water molecules per lipid. All lipids were modeled with the CHARMM36 force-field (23), except Dpop which was modeled using the CHARMM general force field, with atom types determined by the paramchem server. (Gromacs topology file provided in the Supplemental Information.) Initial dimensions in the membrane plane were about 17.5 nm x 17.5 nm, containing approximately 270,000 atoms.

Four different binary mixtures were simulated: Δ9-DOPC: Dpop, Δ11-DOPC: Dpop, Δ9-POPC: Dpop, Δ11 POPC: Dpop. Each binary mixture was simulated at four different compositions: 95:5, 85:15, 70:30, 50:50. Each binary system was simulated for 500 nsec of production simulation as described below. Four additional controls were simulated without any sterol or hopanoid, each for 50 nsec of production simulation as described below: Δ9-DOPC, Δ11-DOPC, Δ9-POPC, Δ11-POPC.

#### Equilibration and production simulations

Each system was prepared individually for production simulation through a series of 6 minimization and heating steps as provided by the CHARMM-GUI equilibration protocol: (i) steepest descent to minimize the initial configuration; (ii) 125,000 steps of leapfrog dynamics with a 1 fsec timestep and velocities reassigned every 500 steps; (iii) 125,000 steps of leapfrog dynamics with a 1 fsec timestep, pressure controlled by the Parinello-Rahman barostat (25) and velocities reassigned every 500 steps, then a total of 750,000 steps of leapfrog dynamics with a 2 fsec timestep and hydrogen positions constrained by LINCS (26), pressure controlled by the Parinello-Rahman barostat (25), and velocities reassigned every 500 steps. During equilibration, double bonds were restrained in the cis configuration to prevent isomerization; these restraints are gradually reduced during the final three stages of the equilibration protocol. Production simulations (NPT ensemble) were integrated with leapfrog using the Parinello-Rahman (25) barostat to control pressure (time constant 5 psec; compressibility 4.5e−5 bar−1; coupled anisotropically to allow independent fluctuation of the in-plane and normal directions) and temperature controlled using Nose-Hoover38,39 (time constant 1 psec) at a temperature of 298K. Hydrogens were constrained with LINCS (expansion order 4), a 2 fsec timestep was used, short range electrostatics were computed directly within 1.2 nm, and long-range electrostatics were computed every timestep using particle mesh Ewald (27, 28) with a grid spacing of 1 Å and cubic interpolation. Long range dispersion was smoothly truncated over 10-12 nm using a force-switch cutoff scheme. Simulations were performed with Gromacs 2020.4.

#### Calculation of simulation observables

The distribution of angles between either Dpop or Chol and the membrane normal was computed, defining the orientation of both by a vector from atom C24 to atom O3. The locations of methyl groups in Dpop or Chol along the direction normal to the membrane were recorded and compiled into histograms with a bin size of 0.87 Å. Deuterium order parameters were obtained from the simulations via

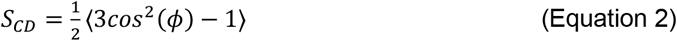

where ϕ is the angle between the C-H bond vectors and the membrane normal at each position along the hydrocarbon chain. The area per lipid was calculated using the Voronoi construction implemented in MEMBPLUGIN (29). The location of each lipid is defined by the center of geometry of the C2, C21, and C31 atoms and the location of the Chol/ Dpop was defined by the O3 atom, and then a Voronoi construction is built around these points in the plane parallel to the membrane surface.

#### Cell culture

*Mesoplasma florum* L1 strains were grown in a modified, lipid-free SP4 media with components as follows (per 1L): Bacto Tryptone 10g, Bacto Peptone 5.3g, PPLO 3.5g, BSA 5.95g, Yeastoleate 2g, D-Glucose 5g, sodium bicarbonate 3.15g, L-Glutamine 0.05g, Penicillin G-sodium salt 0.645g, phenol red 11 mg, pH to 7.0. Lipid diet was added separately prior to passaging at concentration: Dpop, Chol 5 mg/L, egg sphingomyelin 25 mg/L, palmitic acid 10 mg/L, Δ9- and Δ11-PC 12.5 mg/L for the corresponding diets. Cells were grown in glass flasks and incubated at 30°C with shaking at 60 rpm. Growth was recorded using phenol red media pH detection through absorbance at 562nm using a 10mm cuvette (DeNovix DS-11 FX+). Growth rate was defined as negative of the slope of the linearly fitted trendline in the indicative range of phenol red (OD_562nm_ from 0.75 – 0.4).

#### Membrane incorporation

Cells were collected in early exponential stage and centrifuged (5000 rcf, 7 min, 30°C). Supernatant were discarded and cell pellet was washed with wash buffer (200 mM NaCl, 25 mM HEPES, 1% glucose, pH 7.0) and centrifuged (5000 rcf, 7 min, 30°C). The collected pellet was then subjected to a Bligh Dyer extraction (30). Briefly, the pellet was homogenized in a mixture of water:chloroform:methanol in 0.8:1:2 ratio and sonicated for 2 minutes. Subsequently, water and chloroform were added in 1:1 ratio. The mixture was centrifuged at 2000 rcf for 30 seconds in a mini centrifuge to promote phase separation. The lower, organic fraction containing lipids was collected and transferred to a fresh tube. The total lipid extract was then deposited on a silica gel plate (Supelco) and placed in a glass chamber. Chromatography was performed using chloroform as the running phase. After the run, the plate was dried and stained using 8 % copper sulphate in 3% phosphoric acid solution and heated until visible bands were observed. Images were captured using a GelDoc (Biozym Azure c600) and analysed using ImageJ.

#### Membrane osmotic shock

Cells were collected in early exponential stage and centrifuged (5000 rcf, 7 min, 30°C). The collected pellet was resuspended in a serial dilution of 0%, 20%, 40%, 60%, 80% and 100% of wash buffer (200 mM NaCl, 25 mM HEPES, 1% glucose, pH 7.0). The suspension was stained with 10 uM propidium iodide and added to a 96-well plate. Fluoresence emission was recorded using a Tecan Spark fluorescence reader, with excitation at 529 – 549 nm and emission at 609 – 629 nm. The fraction of cell lysed was calculated by normalizing the signal of each sample to the 0% and 100% wash buffer sample. Figure 5C represented the fraction of cell lysed in 80% of wash buffer.

## Notes

### Competing Interest Statement

The authors have declared no competing interest.

### Summary of Updates

Manuscript text shortened for conciseness and clarity. Minor improvements to figures.

## References

1. K. Yamaguchi, M. Kinoshita, Highly stable lipid membranes from archaebacterial extremophiles. Prog. Polym. Sci. 18, 763–804 (1993).

2. O. G. Mouritsen, M. J. Zuckermann, What’s so special about cholesterol? Lipids 39, 1101–1113 (2004).

3. B. J. Belin, et al., Hopanoid lipids: From membranes to plant-bacteria interactions. Nature Reviews Microbiology 16, 304–315 (2018).

4. J. J. Brocks, et al., Biomarker evidence for green and purple sulphur bacteria in a stratified Palaeoproterozoic sea. Nature 437, 866–870 (2005).

5. S. Racolta, P. B. Juhl, D. Sirim, J. Pleiss, The triterpene cyclase protein family: A systematic analysis. Proteins: Structure, Function, and Bioinformatics 80, 2009–2019 (2012).

6. J. P. Sáenz, et al., Hopanoids as functional analogues of cholesterol in bacterial membranes. Proceedings of the National Academy of Sciences of the United States of America 112, 11971–11976 (2015).

7. J. P. Sáenz, E. Sezgin, P. Schwille, K. Simons, Functional convergence of hopanoids and sterols in membrane ordering. Proceedings of the National Academy of Sciences of the United States of America 109, 14236–14240 (2012).

8. H. Martinez-Seara, et al., Interplay of unsaturated phospholipids and cholesterol in membranes: Effect of the double-bond position. Biophysical Journal 95, 3295–3305 (2008).

9. V. Baby, et al., Inferring the Minimal Genome of Mesoplasma florum by. mSystems 3, 1–14 (2018).

10. J. H. Lorent, et al., Plasma membranes are asymmetric in lipid unsaturation, packing and protein shape. Nat Chem Biol 16, 644–652 (2020).

11. G. Chwastek, et al., Principles of Membrane Adaptation Revealed through Environmentally Induced Bacterial Lipidome Remodeling. Cell Reports 32 (2020).

12. J. A. Urbina, et al., Molecular order and dynamics of phosphatidylcholine bilayer membranes in the presence of cholesterol, ergosterol and lanosterol: a comparative study using 2H-, 13C- and 31P-NMR spectroscopy. Biochimica et Biophysica Acta (BBA) - Biomembranes 1238, 163–176 (1995).

13. A. M. Smondyrev, M. L. Berkowitz, Molecular Dynamics Simulation of the Structure of Dimyristoylphosphatidylcholine Bilayers with Cholesterol, Ergosterol, and Lanosterol. Biophysical Journal 80, 1649–1658 (2001).

14. K. Sabatini, J.-P. Mattila, P. K. J. Kinnunen, Interfacial Behavior of Cholesterol, Ergosterol, and Lanosterol in Mixtures with DPPC and DMPC. Biophysical Journal 95, 2340–2355 (2008).

15. E. M. Tookmanian, B. J. Belin, J. P. Sáenz, D. K. Newman, The role of hopanoids in fortifying rhizobia against a changing climate. Environmental Microbiology 23, 2906–2918 (2021).

16. E. Tookmanian, et al., Hopanoids Confer Robustness to Physicochemical Variability in the Niche of the Plant Symbiont Bradyrhizobium diazoefficiens. Journal of Bacteriology 204, e00442–21 (2022).

17. A. J. Fulco, Fatty acid metabolism in bacteria. Progress in Lipid Research 22, 133–160 (1983).

18. M. Makarova, et al., Delineating the Rules for Structural Adaptation of Membrane-Associated Proteins to Evolutionary Changes in Membrane Lipidome. Curr Biol 30, 367–380.e8 (2020).

19. J. Bouwknegt, et al., A squalene–hopene cyclase in Schizosaccharomyces japonicus represents a eukaryotic adaptation to sterol-limited anaerobic environments. Proceedings of the National Academy of Sciences 118, e2105225118 (2021).

20. M. Grzybek, J. Kubiak, A. \Lach, M. Przyby\lo, A. F. Sikorski, A raft-associated species of phosphatidylethanolamine interacts with cholesterol comparably to sphingomyelin. A Langmuir-Blodgett monolayer study. PLoS ONE 4 (2009).

21. S. Jo, T. Kim, V. G. Iyer, W. Im, CHARMM-GUI: A web-based graphical user interface for CHARMM. Journal of Computational Chemistry 29, 1859–1865 (2008).

22. , CHARMM-GUI Input Generator for NAMD, GROMACS, AMBER, OpenMM, and CHARMM/OpenMM Simulations Using the CHARMM36 Additive Force Field | Journal of Chemical Theory and Computation (August 29, 2023).

23. E. L. Wu, et al., CHARMM-GUI Membrane Builder toward realistic biological membrane simulations. Journal of Computational Chemistry 35, 1997–2004 (2014).

24. D. J. Price, C. L. Brooks, A modified TIP3P water potential for simulation with Ewald summation. J Chem Phys 121, 10096–10103 (2004).

25. M. Parrinello, A. Rahman, Polymorphic transitions in single crystals: A new molecular dynamics method. Journal of Applied Physics 52, 7182–7190 (1981).

26. , LINCS: A linear constraint solver for molecular simulations - Hess - 1997 - Journal of Computational Chemistry - Wiley Online Library (August 29, 2023).

27. T. Darden, D. York, L. Pedersen, Particle mesh Ewald: An N⋅log(N) method for Ewald sums in large systems. The Journal of Chemical Physics 98, 10089–10092 (1993).

28. , A smooth particle mesh Ewald method | The Journal of Chemical Physics | AIP Publishing (August 29, 2023).

29. R. Guixà-González, et al., MEMBPLUGIN: studying membrane complexity in VMD. Bioinformatics 30, 1478–1480 (2014).

30. E. G. Bligh, W. J. Dyer, A RAPID METHOD OF TOTAL LIPID EXTRACTION AND PURIFICATION.

