## Supplementary Figures for "Varying the position of phospholipid acyl chain unsaturation modulates hopanoid and sterol ordering" (from another version of this preprint: version 1, 2023-09-06)

### Supplemental Data

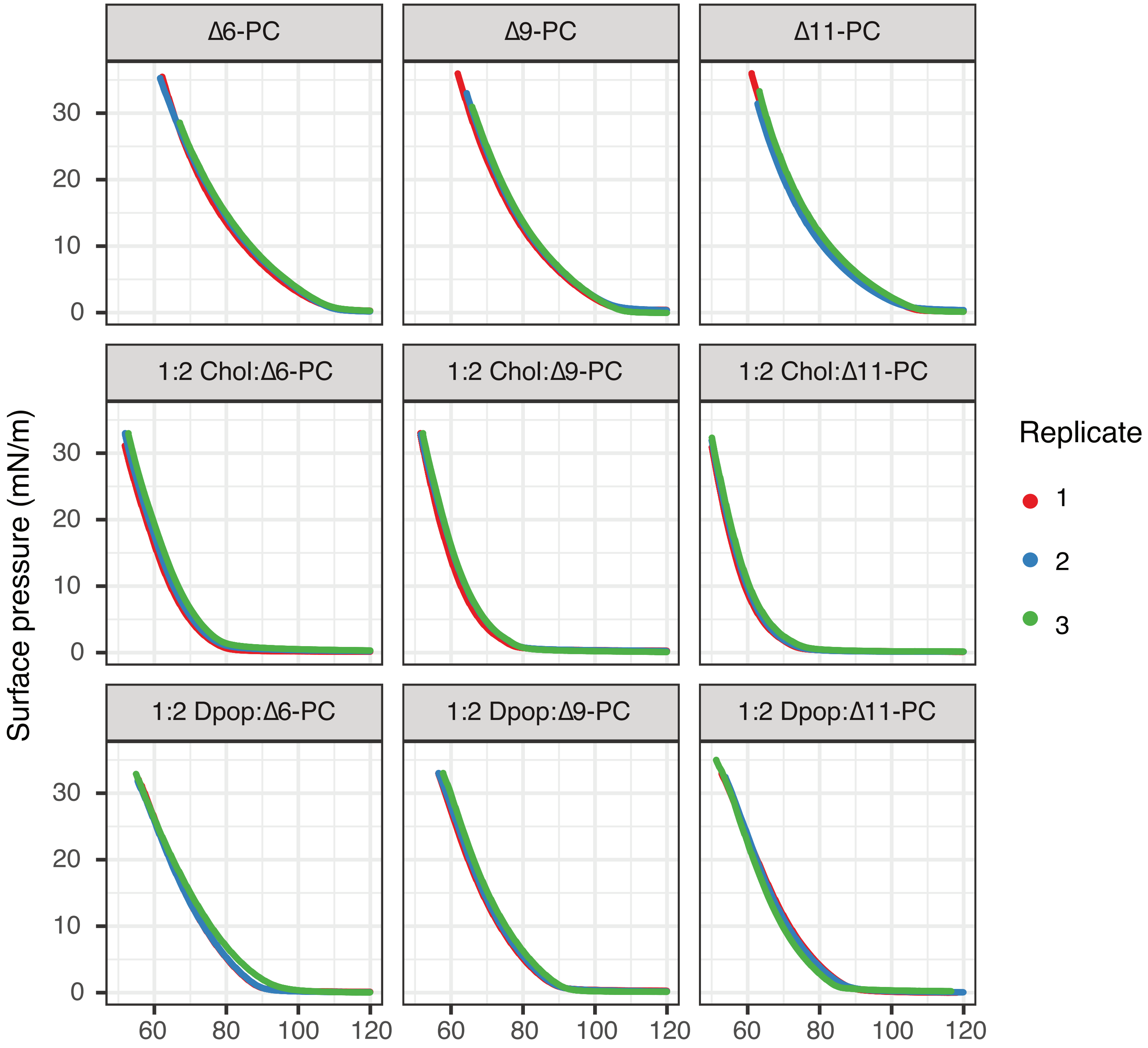


**Fig S1. Raw data of monolayers isotherm replicates.**

The isotherms were fitted to regression for statistical analysis with the following formular:

$$\text{Surface pressure} = \text{coef1} \times \frac{1}{\text{Area}^2} + \text{coef2}$$

**Table S1. Coefficients calculated for each isotherm**

| Sample | Pure | | 1:2 Chol:PC | | 1:2 Dpop:PC | |
|---|---|---|---|---|---|---|
| | coef1 | coef2 | coef1 | coef2 | coef1 | coef2 |
| Δ6-PC | 8.93E-05 | 4.87E-06 | 1.77E-04 | 6.25E-06 | 1.24E-04 | 5.91E-06 |
| Δ6-PC | 8.32E-05 | 5.07E-06 | 1.73E-04 | 6.02E-06 | 1.22E-04 | 6.10E-06 |
| Δ6-PC | 8.33E-05 | 4.89E-06 | 1.66E-04 | 5.85E-06 | 1.09E-04 | 6.49E-06 |
| Δ9-PC | 9.65E-05 | 4.64E-06 | 1.95E-04 | 5.67E-06 | 1.29E-04 | 5.47E-06 |
| Δ9-PC | 9.51E-05 | 4.50E-06 | 1.84E-04 | 5.84E-06 | 1.27E-04 | 5.43E-06 |
| Δ9-PC | 9.44E-05 | 4.43E-06 | 1.85E-04 | 5.58E-06 | 1.25E-04 | 5.16E-06 |
| Δ11-PC | 1.01E-04 | 4.75E-06 | 2.25E-04 | 5.75E-06 | 1.27E-04 | 6.62E-06 |
| Δ11-PC | 1.02E-04 | 4.99E-06 | 2.19E-04 | 5.77E-06 | 1.34E-04 | 6.24E-06 |
| Δ11-PC | 9.70E-05 | 4.77E-06 | 2.12E-04 | 5.88E-06 | 1.40E-04 | 6.37E-06 |

**Fig S2**. Structures of cholesterol (left) and diplopterol (right), with atoms used to define molecular orientation labeled.

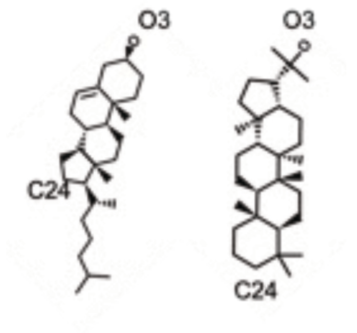


**Fig S3**. Example growth curve through OD562nm read-out. The slope of the curve from 0.75 to 0.4 was used to calculate the growth rate (a.u.)

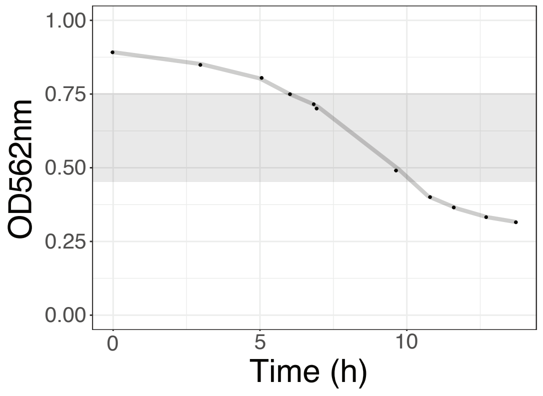


**Fig S4**. Fraction of cell lysed across the osmotic shock gradient tested.

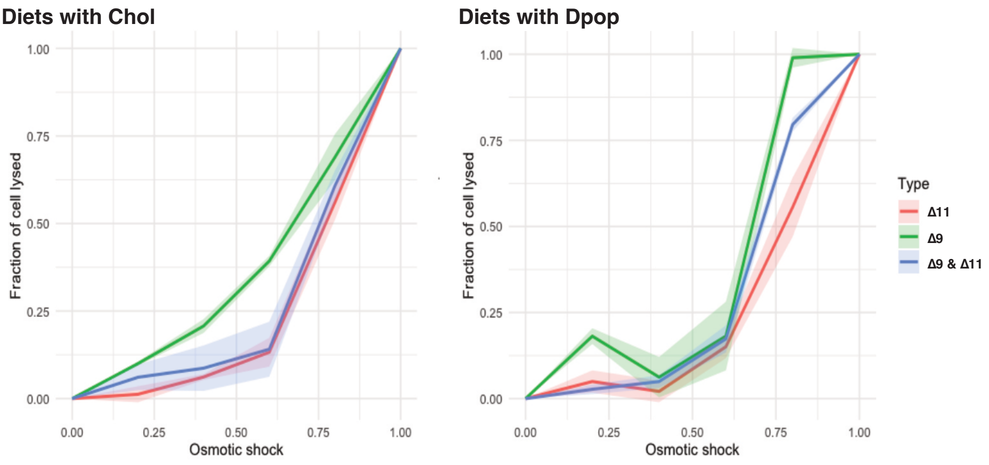
